# An inability to maintain the ribonucleoprotein genomic structure is responsible for host detection of negative-sense RNA viruses

**DOI:** 10.1101/2020.03.12.989319

**Authors:** Daniel Blanco-Melo, Benjamin E. Nilsson-Payant, Skyler Uhl, Beatriz Escudero-Pèrez, Silke Olschewski, Patricia Thibault, Maryline Panis, Maria Rosenthal, César Muñoz-Fontela, Benhur Lee, Benjamin R. tenOever

## Abstract

Cellular biology has a uniformity not shared amongst viruses. This is perhaps best exemplified by negative-sense RNA viruses that encode their genetic material as a ribonucleoprotein complex composed of genome, RNA-dependent RNA polymerase, and the nucleoprotein. Here we demonstrate that limiting nucleoprotein availability not only universally culminates in a replicative catastrophe for negative-sense RNA viruses, but it results in the production of aberrant genomic material and induction of the interferon-based host defenses. This dynamic illustrates the tremendous stress imposed on negative-sense RNA viruses during replication as genomic products accumulate in an environment that requires an increasing demand on nucleoprotein availability. We show that limiting NP by RNA interference or drug targeting blocks replication and primes neighboring cells through the production of interferon. Together, these results demonstrate that the nucleoprotein represents the Achilles heel of the entire phylum of negative-sense RNA viruses. Here we establish this principle for a diverse collection of human pathogens and propose that the nucleoprotein should be a primary target for the development of future antiviral drugs.

**HIGHLIGHTS:**

- Limited levels of NP result in production of defective viral genomes
- Defective viral genomes and viral antagonists are key determinants of the host antiviral response
- The host response and defective viral genome generation further exasperate NP availability
- NP is an optimal drug target for the whole phylum of negative-sense RNA viruses

## INTRODUCTION

Negative-sense RNA viruses comprise many of the most significant pathogens that account for both human and veterinary viral diseases. These viruses have shaped the history of humankind and are a significant burden on global health. Over the past 150 years, measles and influenza A virus (IAV) alone have been estimated to have killed over 300 million people worldwide (Goodson and Seward, 2015). While vaccines have significantly diminished these numbers, measles virus continues to cause annual morbidity and mortality amongst unvaccinated populations (Clemmons et al., 2017). IAV also continues to cause a global health burden because seasonal strains genetically drift to render vaccines ineffective (Lowen, 2017). Moreover, IAV also has the capacity to re-assort with other IAV strains, a process referred to as antigenic shift, which can result in a virus capable of rapidly spreading to pandemic proportions (Lowen, 2017). In addition to these known agents for which therapeutics have been developed, countless novel viral pathogens that lack widely available vaccines, such as Nipah, Ebola, and Lassa viruses, continue to emerge as the human population expands into uncharted microbe-laden areas (Plowright et al., 2017).

The notoriety of negative-sense RNA viruses can be attributed, in part, to their unique biology. All life, regardless of domain or kingdom, store and process information essentially in the same manner. Genomic DNA represents the long-term storage of information that is processed as RNA. RNA serves both functional roles and as an intermediate for protein production which together assemble into all of the components necessary for cellular life. Although this flow of biological information is universal in all living things, viruses have evolved ways to rewrite these dynamics in some cases avoid the need for a DNA intermediate at all. One of the more unique strategies for replacing DNA as the long-term storage of information derives from that of negative-sense RNA viruses that maintain their genomes in an antisense orientation bound by viral nucleoprotein (NP) and the RNA-dependent RNA polymerase (RdRp) (Lou, 2018).

As a result of storing their genomic information in a manner that deviates from the central dogma, these viruses can be easily recognized by the host cell as ‘non-self’. While the orientation of the RNA is not considered to be a pathogen-associated molecular pattern (PAMP), these genomic RNAs are generally unmodified on their 5’ end and therefore begin with a 5’ tri-phosphate motif that is a substrate for the pattern recognition receptor (PRR), RIG-I (Gebhardt et al., 2017). In addition to this PAMP, RNA viruses are generally also thought to generate a significant amount of double stranded RNA which can be recognized by the PRR, MDA5 (Schneider et al., 2014). In both examples, these PAMPs can derive from both full length genomic products or defective viral genomes (DVGs) which are known to accumulate to high numbers during the replicative process (Vignuzzi and Lopez, 2019). Following PAMP recognition, PRR induces its oligomerization and the assembly of a variety of scaffold proteins and kinases (Schneider et al., 2014). These complexes orchestrate the activation of both NFκB and IRF transcription factor families and the subsequent induction of antiviral cytokines including type I and III interferons (IFN-I and IFN-III) (Levy et al., 2011). IFN-I and -III signal in both an autocrine and paracrine manner to induce hundreds of interferon stimulated genes (ISGs), such as IFIT1 (IFN-induced proteins with tetratricopeptide repeats), which can block the translation of RNA with unusual 5’ ends (Kumar et al., 2014). The collective induction of ISGs serves to block many aspects of cell biology as a means of slowing virus infection and provide time for the adaptive immune response to ultimately clear the pathogen (Iwasaki and Medzhitov, 2015). Given the effectiveness of the IFN-based defenses, negative-sense RNA viruses generally encode at least one antagonist of this host antiviral system (Garcia-Sastre, 2017). In this work we describe a critical balance between NP and negative-sense RNA viral products which, if disrupted leads to the accumulation and recognition of aberrant viral genomes that overwhelm the viral antagonist and result in the induction of the host antiviral response.

## RESULTS

### Characterizing negative-sense RNA virus replication under limiting NP levels

Given the knowledge that negative-sense RNA viruses are generally sensed through the recognition of genomic-based PAMPs, we sought to determine how the levels of NP may influence this dynamic. To this end, we tested the robustness of the viral ribonucleoprotein (RNP) complex to NP perturbations, and investigated their effects in the production and recognition of PAMPs. In order to generate an imbalance in NP production, we engineered recombinant strains of IAV and Sendai virus (SeV) in which the NP transcript would be the target of host-derived miRNAs previously shown to exert potent silencing activity in a broad range of cells (Figure S1A-C)(Aguado et al., 2018).

To determine the consequences of limiting NP, we infected lung epithelial cells with miRNA-targeted (herein denoted by NP-T) or untargeted control (denoted NP-C) viruses and evaluated both host and pathogen biology. Targeting of NP mRNA resulted in an inability to detect the NP protein both in the context of SeV and IAV infection (Figure 1A-B). Despite the reduction in replication caused by the loss of NP, miRNA-targeted viruses induced a greater host response than untargeted viruses as illustrated by the induction of IFIT1 (Figure 1A-B). Moreover, RNA sequencing corroborated that miRNA-targeting of NP resulted in a dramatic reduction of viral reads that exceeded multiple orders of magnitude when compared to untargeted control virus (Figure 1C-D). In addition, mRNA-Seq demonstrated that despite the decrease in viral RNA levels, both SeV and IAV infections in which NP was depleted resulted in a robust Type I and III interferon (IFN-I - III) response as well as the induction of PRRs (DDX58, IFIH1, TLR9), transcription factors (IRF1, IRF7, IRF9) and antiviral effector proteins (BST2/Tetherin, IFIT1-3, MX1) (Figure 1E-F and Table S1-S2). The host response to virus infection in which NP was limiting was also more robust, relative to viral load, in both the number of genes induced and their overall induction when compared to control infections (Figure S1D-E and Table S1-S2).

**Figure 1:**
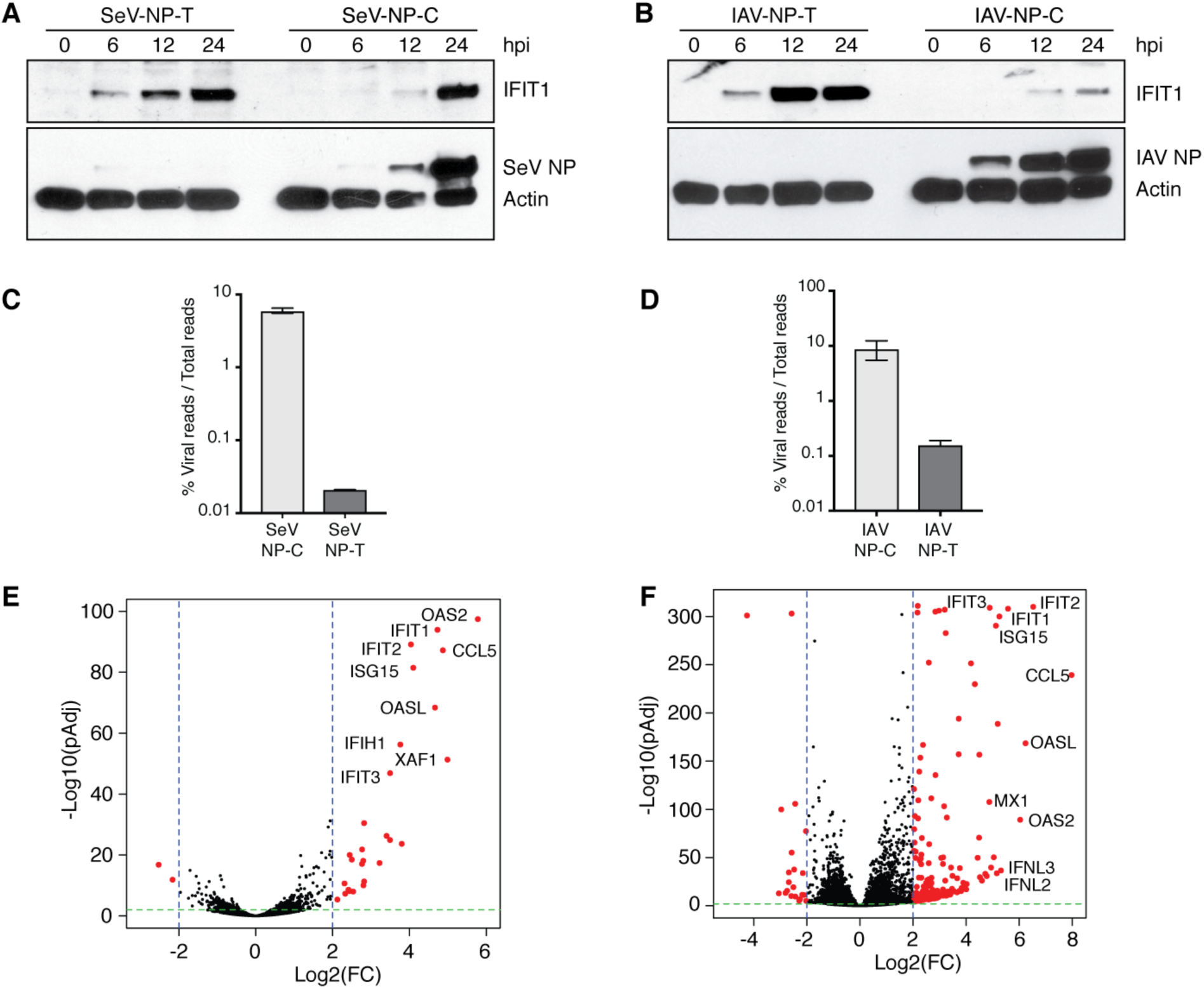
Limiting NP availability severely impairs viral infections and leads to the induction of an antiviral host response. (**A**-**B**) A549 cells were infected either with SeV-NP-T and SeV-NP-C (**A**) or IAV-NP-T and IAV-NP-C (**B**) for 0, 6, 12 and 24 h at an MOI of 5. Whole cell lysates were analyzed by western blot using primary antibodies for human IFIT1, Pan-Actin and either SeV or IAV nucleoprotein (NP). (**C**-**D)** Percentage of NGS viral reads from infected A549 cells with (**c**) SeV-NP-T/C (MOI = 5, 24 hpi) or (**D**) IAV-NP-T/C (MOI = 5, 9 hpi). Values are given as mean ± sd of mRNA-seq reads aligning to corresponding viral genomes over total number of reads (n = 2). * p < 0.05 by an unpaired t test with Welch’s correction; ns: Non significant. **E**-**F**, Volcano plots depicting the host transcriptional response to (**E**) SeV-NP-T and (**F**) IAV-NP-T compared to mock infected cells. Differential expression analyses correspond to samples used in **C**-**D**. Red points represent statistically significant differentially expressed genes (DEGs) in A549 cells infected with the corresponding viruses compared to mock infected cells (Log2(FC) > 2, pAdj < 0.01, n=2).

### NP influences RdRp processivity and DVG levels of negative-sense RNA viruses

To determine the molecular basis for the robust IFN-I response despite near undetectable levels of viral replication, we next focused on the genome integrity of NP-targeted viruses (Figure 2). To this end, we compared viral RNA reads from infections within a single round of replication. Interestingly, when analyzing non-contiguous viral reads spanning genomic junctions from either IAV and SeV, we observe an increase in these reads, denoting defective viral genomes, in conditions in which NP is limiting (Figure 2A and C). Moreover, the alignment profile of NP-targeted IAV demonstrated the greatest enrichment in reads mapping to the ends of segments 1-3, characteristic of DVG formation (Figure 2B) (Baum et al., 2010). While this same trend was less apparent when mapping SeV negative-sense genomic RNA, paramyxoviruses largely induce DVGs comprised of so called ‘copy-back’ genomes which cause a selective increase only in terminal end reads (Patel et al., 2013) (Figure 2D). Moreover, it is worth noting that this measurement is a significant underrepresentation of the amount of DVGs being formed as this analysis is comparing the percent of junction reads to total genomic alignments. Taken together, these data suggest that limiting NP availability induces the formation of DVGs.

**Figure 2:**
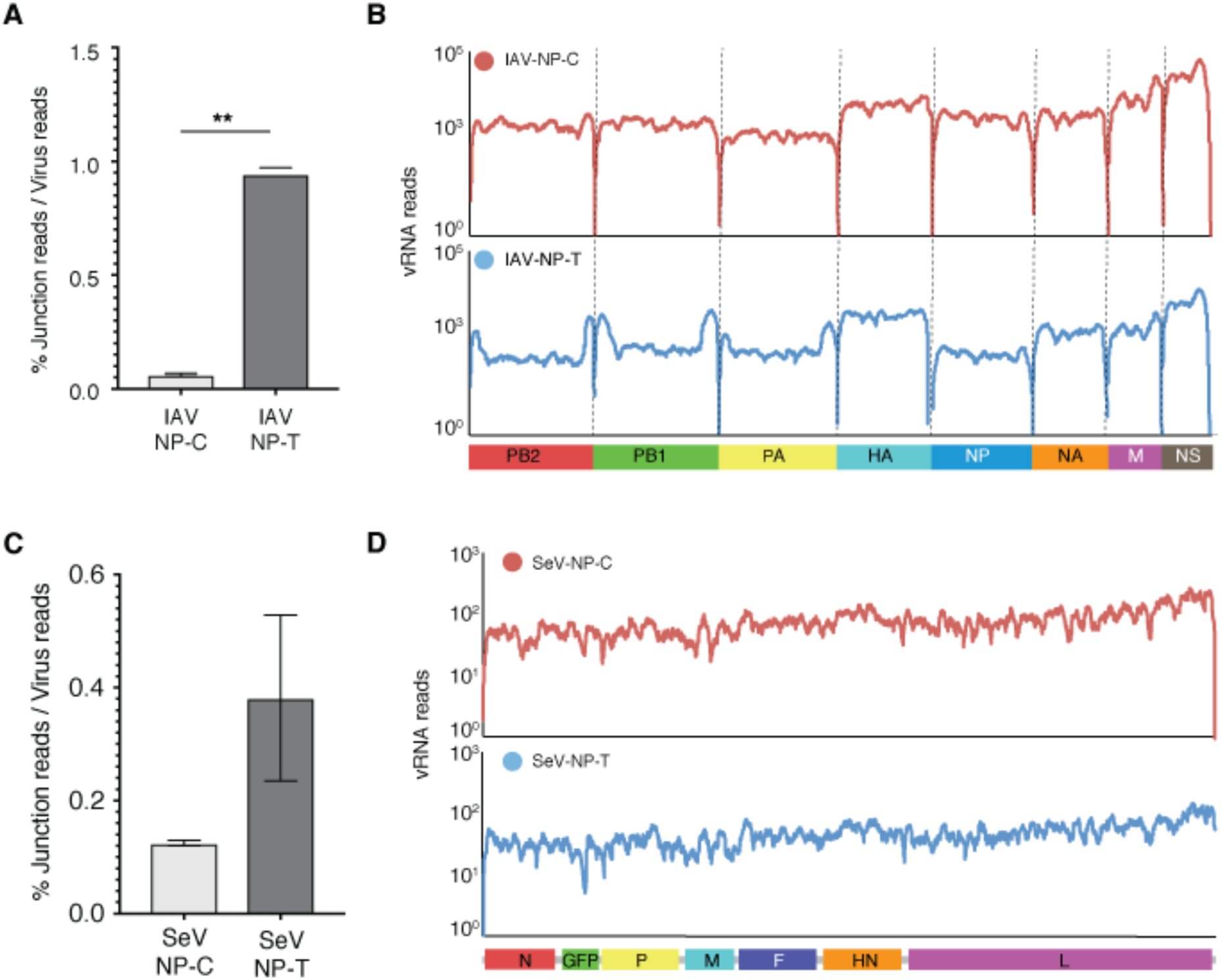
Limiting NP availability results in the accumulation of aberrant vRNAs. (**A**) Proportion of DVGs present in A549 cells infected with IAV-NP-T/C (MOI = 5, 9 hpi). Values are given as mean ± sd of ribosome-depleted total RNA sequencing reads spanning junction sites in corresponding viral genomes over total number of viral reads (n = 2). ** p < 0.01 by an unpaired t test with Welch’s correction. **(B)** Read coverage along IAV-NP-C (top) and IAV-NP-T (bottom) viral genomes in A549 cells infected with the corresponding viruses (Moi = 5, 9 hpi). Results from ribosome-depleted total RNA sequencing reads aligning to corresponding viral genomes. **(C)** As in **A** for SeV-NP-C and SeV-NP-T (MOI=3, 9 hpi). **(D)** As in **B** for SeV-NP-C and SeV-NP-T.

In an effort to better understand the relationship between NP levels and DVG production, we directly analyzed the population of different viral RNAs produced under limiting NP conditions in IAV infections by Northern blot. We focused on IAV largely because the genomic segmentation of the virus allowed us to better control the relative levels of NP, RdRp, and genomic template. To assess the role of NP in DVG production, we first focused on the RNA transcripts that accumulated in response to our recombinant IAV-NP-T and -C strains. To this end, we found that infection with the untargeted NP control virus resulted in the accumulation of full length vRNA and the previously characterized small viral RNA (svRNA) (Figure 3A) (Perez et al., 2010; Perez et al., 2012; Umbach et al., 2010). In contrast, while a complete loss of full length vRNA was observed in the NP-targeted virus, these same conditions showed the accumulation of a range of smaller viral RNA species comparable in size to the recently described mini viral RNA (mvRNA) – an NP-independent DVG (Figure 3A) (Te Velthuis et al., 2018). In agreement with mvRNA being a potent inducer of the antiviral response, we also observed a correlation between its appearance and the induction of IFNβ (Figure 3B). To further confirm that the production of aberrant vRNA genomes and the induction of IFNβ was directly linked to an inefficient amount of NP present during viral RNA genome replication, we transiently expressed a 200 nucleotide-long vRNA template mimicking segment six, together with the three RdRp subunits: PB2, PB1 and PA in the presence of increasing amounts of NP (Figure 3C-D). Consistent with the results in Figure 3A, increasing NP availability resulted in the generation of full-length vRNA (Figure 3C). In conditions where NP was limited however, we observe the production of mvRNA. In contrast, in the complete absence of NP, we observed no viral RNA products for the 200 nt minigenome, consistent with the findings that vRNA greater than 76 nt require NP to replicate (Figure 3C) (Turrell et al., 2013). Taken together, these results suggest that while high levels of NP are required for efficient genomic replication, insufficient NP results in the production of DVGs. Moreover, under these same conditions, we observe low NP levels correlate with the induction of IFNβ (Figure 3D). These data further allow us to conclude that the ISG response to IAV-NP-T is the result of limiting NP, rather than a corresponding loss of NS1, as this same ISG signature is lost in conditions of excess NP in an NS1-independent system (Figure 3B and 3D). These data are also consistent with the observation that when the genome of vesicular stomatitis virus (VSV) is rearranged, there is an observable enhancement in ISGs that correlated to decreased transcription of the nucleoprotein (Pesko et al., 2015).

**Figure 3:**
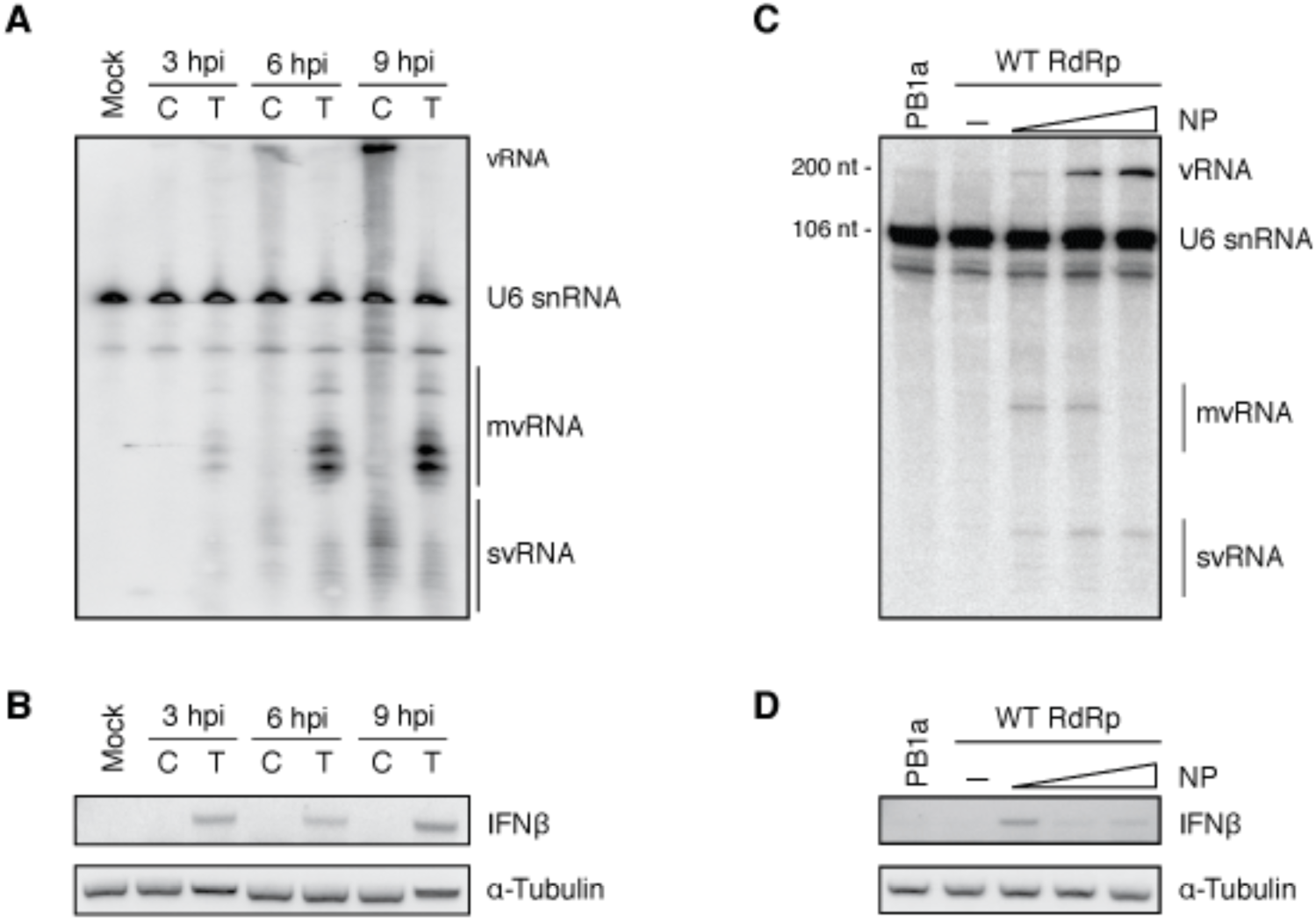
Limiting NP availability results in the production of defective viral genomes. (**A**) A549 cells were infected with IAV-NP-T (T) or IAV-NP-C (C) virus at an MOI of 5 and RNA was extracted 3, 6 and 9 hpi. Total RNA was assessed by Northern blot analysis using radiolabelled probes against human U6 snRNA as well as the conserved 5′ terminus of the negative-sense vRNA promoter of IAV. (**B**) RNA obtained in **A** was analyzed by RT-PCR for the presence of IFNβ and α-Tubulin mRNA transcripts. (**C**) HEK-293T cells were cotransfected with plasmids expressing a mutant 200-nt long IAV segment 6 vRNA (containing an internal deletion leaving only the 100 terminal nucleotides at the 3′ and 5′ vRNA ends), the IAV RdRp subunits PA, PB1 and PB2 and increasing amounts of NP. Accumulation of negative-sense viral RNA was assessed by Northern blot analysis of total extracted RNA using radiolabelled probes against human U6 snRNA as well as the conserved 5′ terminus of the negative-sense vRNA promoter of IAV. (**D**) RNA obtained in **C** was analyzed by RT-PCR for the presence of IFNβ and α-Tubulin mRNA transcripts.

### DVG content dictates the transcriptional response to negative-sense RNA viruses

Given the observation that limiting NP results in the production of DVGs and IFNβ, we next assessed how this biology would impact the overall host response to infection. To this end, we generated stocks of IAV containing low or high levels of DVGs (herein denoted LD and HD, respectively), as well as LD and HD stocks of IAV lacking NS1 (ΔNS1) (Garcia-Sastre et al., 1998) which we could confirm using standard RT-PCR (Figure 4A). Using these four unique populations of IAV, we infected primary human lung epithelial (NHBE) cells and performed RNA-Seq to further verify the relative levels of DVGs (Figure 4B and S2). In addition to aligning virus-derived reads, we also compared the cellular response to each of these infections and contrasted them to mock treatment or cells administered IFNβ (Figure 4C). Compared to mock treated cells, we observed ∼250 genes significantly induced in response to IFNβ (Log_2_(Fold Change) > 2, pAdj < 0.05) which largely comprise the canonical list of known ISGs (Schneider et al., 2014). Of these ISGs, we observed only an approximate 10% overlap when these same cells were infected with wild type virus containing a low level of DVGs (IAV LD) – suggesting that in the absence of DVGs, the transcriptional response is largely independent of IFN-I or IFN-III. In contrast, the same virus containing high DVG (IAV HD) content resulted in ∼40% of the ISG signature – even in the presence of NS1. When comparable conditions were generated in the absence of NS1 (ΔNS1 HD), the host response included over 1500 differentially expressed genes (DEGs) and encompassed the complete breadth of the ISG response (Figure 4C and Table S3-S5). Use of ΔNS1 LD generated a host response that was intermediate of IAV wild type vs. ΔNS1 with high DVG content. Together, these data support the idea that the host response to negative-sense RNA virus infection is defined by the levels of DVGs, by-products of limited NP availability, and that this activity can be buffered with the expression of a virus antagonist such as NS1 in agreement with single-cell analyses (Russell et al., 2019).

**Figure 4:**
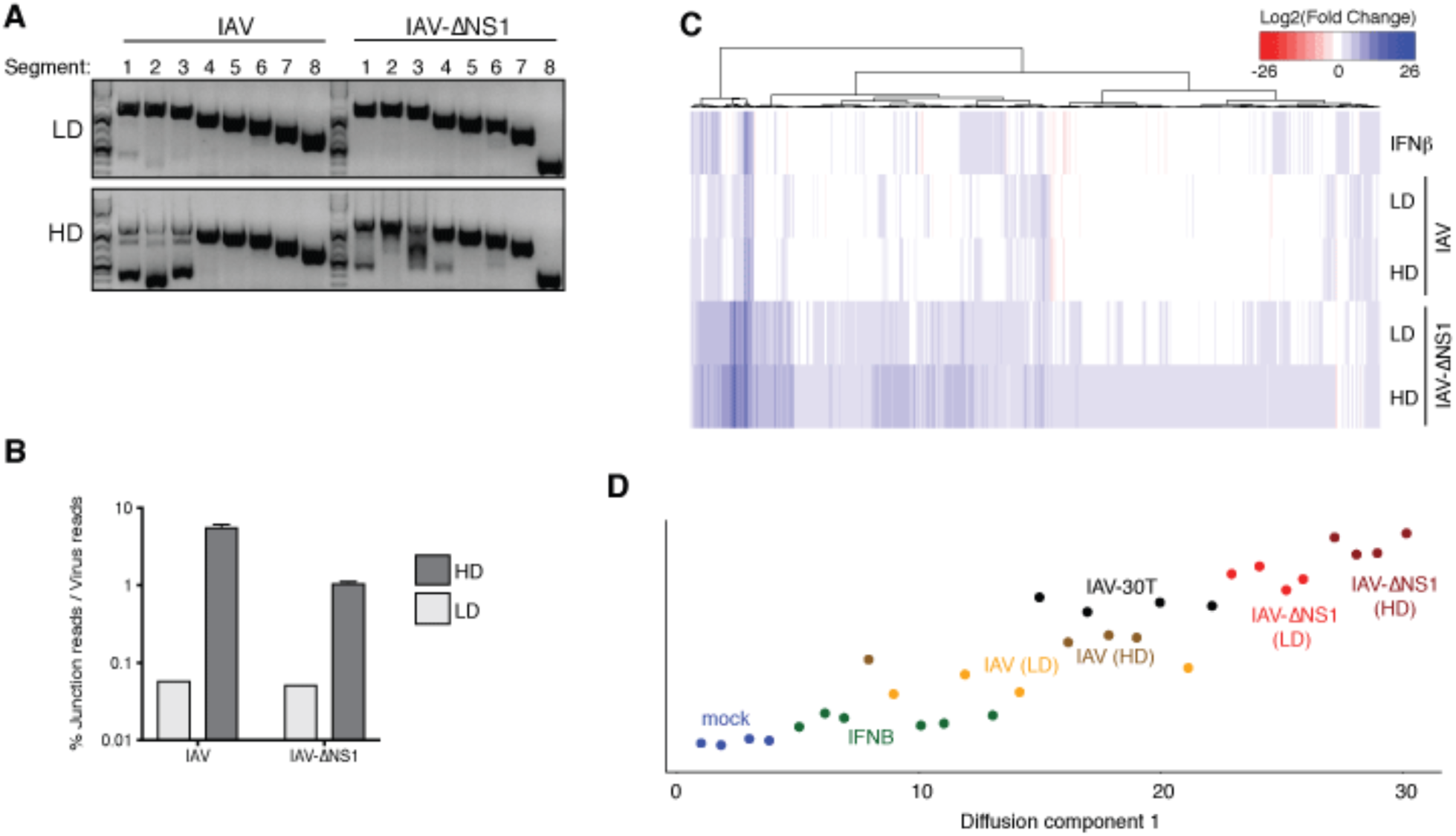
The host response to IAV is defined by DVG content and NS1 expression. (**A**) RNA prepared from virus stocks with low (LD) and high (HD) DVG content was reverse transcribed using a universal terminal 5′ vRNA primer prior to segment specific PCR. Size markers on the left are given in nucleotides. (**B**) Analysis of DVGs present in NHBE cells infected with LD or HD versions of wild-type IAV or IAV ΔNS1 virus (MOI = 3, 12 hpi). Values are given as mean ± sd of mRNA-seq reads spanning junction sites in corresponding viral genomes over total number of viral reads (n = 3). (**C**) Heatmap depicting the transcriptional response of NHBE cells to IFNB treatment or infection with LD or HD versions of IAV WT or IAV ΔNS1 virus (n = 4, MOI = 3, 12 hpi). Values represent the Log_2_(fold change) expression (compared to mock-treated cells) for statistically significant DEGs in at least one condition. Values for IFN treatment represent an aggregate of cells treated for 4, 6 and 12 hrs compared to mock-treated cells (n = 2 for each time point). (**D**) Diffusion map of the transcriptome of NHBE cells upon IFNB treatment or infection with different IAVs. Graph depicts samples ordered only by diffusion component 1 (n=4). IAV-30T: NS1-T targeted IAV. LD: IAV with low DVG content. HD: IAV with high DVG content.

To further understand the role of NS1 as a buffer to mask DVGs, we utilized a previously described recombinant IAV in which the NS1 and NEP open reading frames are separated to enable the addition of two miRNA targets that silence NS1 while sparing NEP (Figure S3A)(Chua et al., 2013). To achieve intermediate levels of the antagonist in NHBE cells, two target sites to miR-30 were incorporated (IAV-NS1-T), as this miRNA is moderately expressed in NHBE cells (McCall et al., 2017) (Figure S3B). Infection in MDCK-NS1 and NHBE cells showed that the incorporation of miR-30 targets into segment eight successfully reduced NS1 protein to levels intermediate between ΔNS1 and wild type virus (Figure S3C-D). Given the success in reducing NS1 in NHBE cells, we next performed RNA-Seq using these viral recombinants.

To correlate the transcriptional response to infection in an environment with limited NS1, we constructed a diffusion map to compare all of the transcriptional profiles in NHBE cells performed thus far (Figure 4D). Using this type of multidimensional reduction analyses we observe that biological replicates of infected NHBE cells clustered tightly together following a progression of transcriptional changes between samples. That is, in comparison to mock, the transcriptional footprint increases with the addition of IFNβ followed by IAV infections which are further delineated by the balance of DVG content and NS1 levels (Figure 4D). For example, Along the first diffusion component IAV LD (low DVG, high NS1) is left of IAV HD (high DVG, high NS1), which maps closely to the administration of IFNβ alone. To the right of these samples falls IAV ΔNS1 LD (low DVG, no NS1) followed by the greatest transcriptional response observed with IAV ΔNS1 HD (high DVG, no NS1). Consistent with the idea that NS1 buffers the host response to DVG content that can arise as a result of limited NP, we find IAV-NS1-T (high DVG, moderate NS1) is placed intermediate to IAV HD and IAV ΔNS1 LD (Figure 4D and Table S3-S6). In summary, we demonstrate here that the host transcriptional response is directly correlated to the levels of DVGs as well as NS1 in the cell.

### Defining the priorities of NS1 antagonism on the host response to infection

As NS1 buffers the response to DVG production, we next determined how limiting NS1 impacts the host response as a proxy for defining its antagonistic potency. That is, DEGs between IAV-NS1-T and IAV ΔNS1 would indicate host responses that can be blocked with only moderate levels of NS1 (Figure 5A and Table S7-S8). In contrast, host responses that are comparable between IAV-NS1-T and IAV ΔNS1 as compared to wild type would indicate pathways that require a high NS1 pool to achieve inhibition (Figure 5B and Table S7-S8). Based on this logic, we first examined genes in which IAV-WT and IAV-NS1-T were comparable and distinct from IAV ΔNS1 (Figure 5A). Enriched biological processes from this subset of host transcripts included an array of cytokine signaling pathways and inflammatory responses denoted in part by the inclusion of: IFNβ, CCL20, CCL4, and IL1A (Figure 5C and Table S8). However, when comparing DEGs between wild type virus and conditions in which NS1 was absent or limiting, we observed the expression of chaperones involved in the response to unfolded proteins in addition to a number of ISGs (Figure 5B, 5D, and Table S8). These results could be further confirmed by RNA-Seq data of IAV ΔNS1 infections of NHBE cells that demonstrate a comparable cascade of host transcription defined by the time of infection (Figure S4 and Table S9).

**Figure 5:**
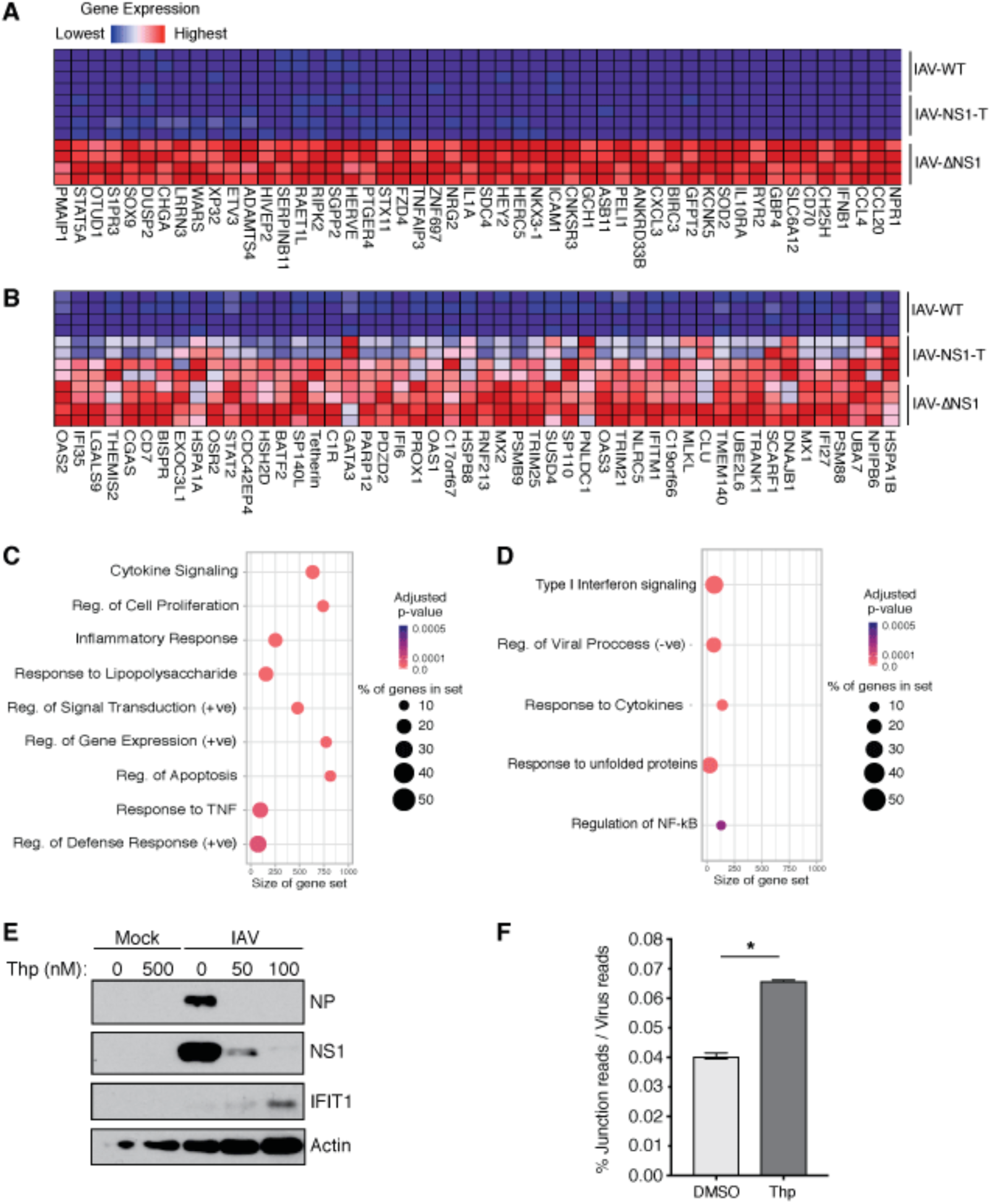
NS1 prioritization of antagonizing the host antiviral response. (**A-B**) Heatmaps depicting the gene expression of the top 50 host gene markers (defined by gene set enrichment analyses) in NHBE cells infected with WT or recombinant IAVs. (**A**) NS1 negative vs. NS1 positive IAVs. (**B**) WT vs. mutant NS1 IAVs. (**C-D**) Top enriched GO terms in cells infected with: (**C**) NS1 negative IAV compared to NS1 positive IAVs or (**D**) WT IAV compared to mutant NS1 IAVs. (**E**) A549 cells were infected with IAV in the presence of the indicated concentrations of thapsigargin. Whole cell lysates were analyzed by Western blot. **(F)** Proportion of DVGs present in NHBE cells infected with IAV ΔNS1 (MOI = 10) and treated with 1 μM of thapsigargin for 12 hrs. Values are given as mean ± sd of ribosome-depleted total RNA sequencing reads spanning junction sites in corresponding viral genomes over total number of viral reads (n = 2). * p < 0.05 by an unpaired t test with Welch’s correction.

As both the response to unfolded protein and IFN-I result in shutdown of the host translational machinery through PERK and PKR, respectively, this would suggest that induction of either of these pathways would result in cell-mediated limiting of viral proteins (Baird and Wek, 2012; Garcia et al., 2007). As negative-sense RNA viruses demand more NP than any other viral transcript, host cell targeting of translation would result in DVG production prior to replicative shutdown. This dynamic would generate a feed forward loop in which aberrant viral RNA by-products of replication lead to shutdown of translation, and the subsequent generation of more aberrant viral RNA ultimately resulting in viral clearance or the induction of cell death. To formally investigate how the antiviral pathways that target translation would impact IAV biology, we infected cells treated with increasing amounts of thapsigargin, a compound that induces the unfolded protein response (UPR) by inhibiting the function of calcium-dependent chaperones (Figure 5E) (Sehgal et al., 2017). Consistent with our previous results, these data demonstrated that engagement of the UPR rapidly resulted in loss of NP and a corresponding induction of the host response (as measured by IFNβ and IFIT1). Moreover, treatment with thapsigargin also correlated with a significant increase in DVG production (Figure 5F).

### Targeting NP shows universal engagement of the host response independent of negative-sense RNA viral family

Given that the host response was largely defined by DVG content, we next chose to determine whether targeting of NP would provide greater protection than another essential viral gene. To this end, we treated cells with siRNAs targeting IAV NP or PB1 transcripts prior to infection with IAV-ΔNS1 to enable the full breadth of the cellular antiviral response (Figure S5A). In agreement with our hypothesis, this experimental framework demonstrated that loss of NP resulted in an increase in IFIT1 induction as compared to either scrambled or polymerase-based siRNAs. Moreover, when we compared NP siRNAs to scrambled control siRNAs in wild type and STAT1-deficient cells, we could corroborate IFIT1 induction in conditions where NP is limiting and showed that this required intact IFN-I signaling as IFN-independent IFIT1 expression correlated with virus replication (Figure S5B).

To further determine whether targeting of NP could enhance the antiviral host response for other negative-sense RNA viruses, we infected siRNA-treated cells with a wide range of human pathogens including BSL2, BSL3, and BSL4 pathogens (i.e. IAV, human parainfluenza virus 3 (HPIV3), measles virus (MeV), human respiratory syncytial virus (HRSV), VSV, Ebola virus (EBOV), and Lassa Fever virus (LASV)) (Figure 6A-G). In each of these examples, limiting NP was found to effectively block virus replication while still potently engaging IFN-I and the host antiviral response as observed by the induction of IFIT1. Furthermore, as expected based on Western blot analysis of whole cell lysates in Figures 6A-G, limiting NP also efficiently reduced the number of infected cells (Figure S6A) or demonstrated significant inhibition of viral replication and therefore reduced reporter gene expression across all viruses tested (Figure S6B).

**Figure 6:**
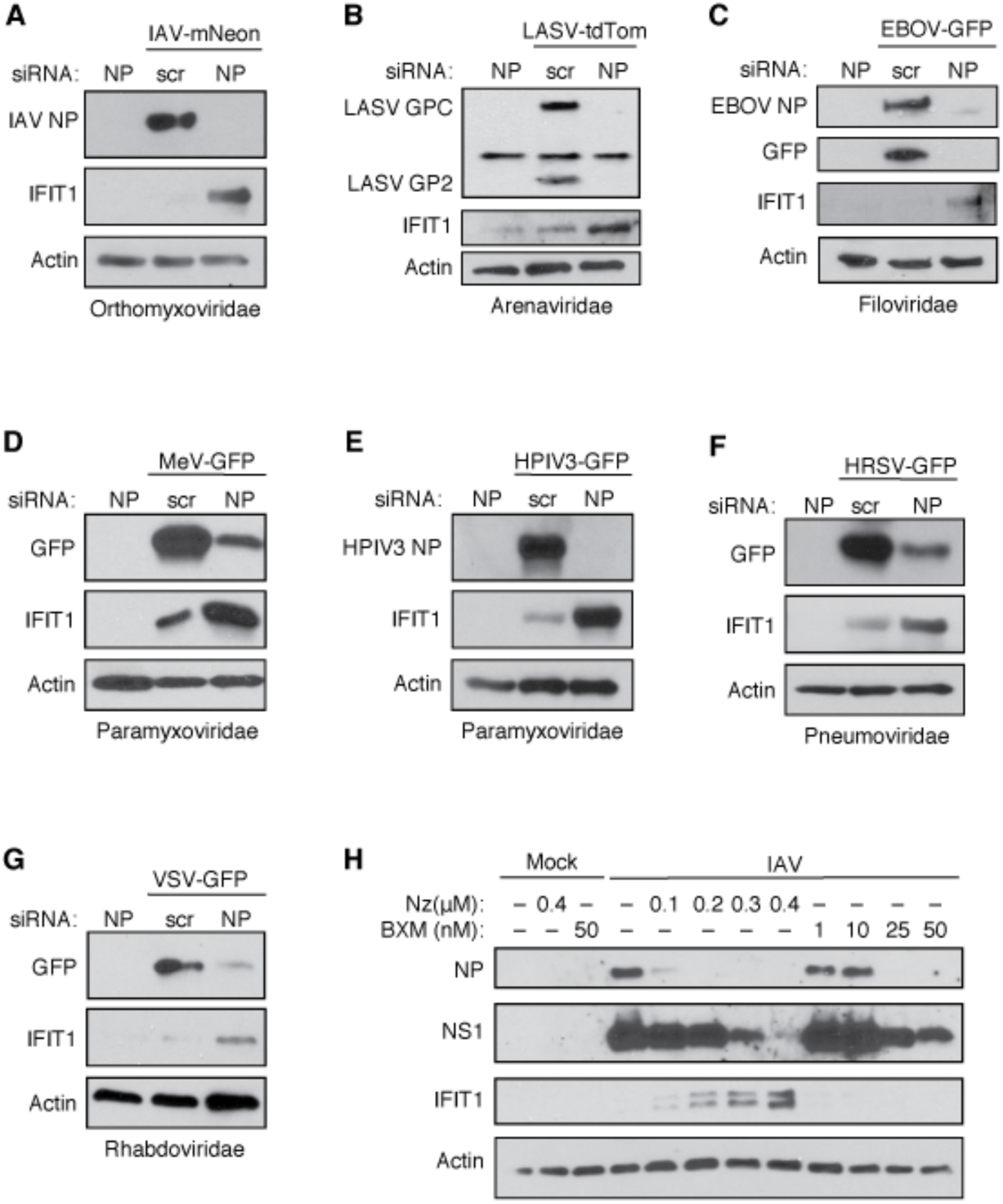
NP as the “Achilles heel” of negative-sense RNA viruses. (**A-G**) A549 cells were transfected with siRNA pools targeting virus specific NP or scrambled controls (scr) 24hrs prior to infection with the indicated viruses. Whole cell lysates were analyzed by Western blot using primary antibodies against human IFIT1, Pan-Actin as well as a viral product as indicated. (**A**) Influenza A Virus (IAV), (**B**) Lassa virus (LASV), (**C**) Ebola virus (EBOV), (**D**) Measles (MeV), (**E**) Human Parainfluenza Virus 3 (HPIV3), (**F**) human respiratory syncytial virus (HRSV), and (**G**) Vesicular Stomatitis Virus (VSV). (**H**) A549 cells were infected with IAV for 8 h at an MOI of 5 in the presence of the indicated concentrations of the known IAV nucleoprotein inhibitor nucleozin (Nz) and RdRp inhibitor Baloxavir marboxil (BXM). Whole cell lysates were analysed by Western blot using primary antibodies against human IFIT1, Pan-Actin as well as the viral NP, and NS1 proteins.

### Drug targeting of NP demonstrates bystander effect

To determine if it was the silencing of NP that specifically resulted in the production of DVGs and the induction of IFNβ, we next tested two different small molecule inhibitors of IAV targeting either NP (Nucleozin, Nz) or PA (Baloxavir marboxil, BXM)(Hayden et al., 2018; Kao et al., 2010). Nz is thought to cause the aggregation of NP in IAV infection, thus preventing nuclear accumulation and normal NP function in replication and transcription of the viral RNA genome (Kao et al., 2010). BXM is an inhibitor of the endonuclease domain of the PA subunit of the viral RdRp, therefore directly inhibiting viral mRNA synthesis (Hayden et al., 2018). Using increasing concentrations of Nz or BXM, we found that we could effectively block replication with both inhibitors in a concentration dependent manner (Figure 6H). However, despite the effectiveness of both inhibitors, only the targeting of NP with Nz resulted in the induction of IFIT1. These results, in agreement with all of the data presented herein, demonstrate that specifically targeting NP leads to the successful engagement of the cellular antiviral defenses thereby providing an additional bystander level of protection.

## DISCUSSION

The data presented in this work provide a comprehensive understanding of the events that culminate in IFNβ induction following the replication of negative-sense RNA viruses. Following entry and transcription, the accumulation of different viral products and a changing cellular environment induces a switch to *de novo* genome production. This process initiates an essential activity for virus replication but also one that will ultimately result in detection by the host. As new genomes are generated, there is an immediate demand for available NP pools. As this dynamic progresses, NP inevitably becomes limiting thereby forming only a partial RNP complex, which promotes the formation of DVGs, perhaps in part by providing the viral RdRp with more flexibility to slide or disengage from the template strand during replication (Te Velthuis et al., 2018). DVGs generated by this process further exacerbate the problem as they too can be substrates for NP and the polymerase, thereby overwhelming the resources of the virus. Given the space in which negative-sense RNA viruses must operate, generating an environment in which RNA substrate exceeds available NP pools would be unavoidable. This is exemplified by the fact that even doses well below the reported EC50 of nucleozin can effectively induce replication catastrophe (Kao et al., 2010).

As replication must proceed even in the face of suboptimal NP levels, negative-sense RNA viruses require potent antagonists of the antiviral host defenses to ameliorate the consequences of recognition. This dynamic is likely also true in the context of negative-sense RNA virus infection in arthropods and/or plants were RNAi is the dominant antiviral defense system (tenOever, 2016). In this arena, DVG production has been shown to serve as a substrate for the generation of siRNAs (Poirier et al., 2018). For this reason, many antagonists such as NS1 have been found to interfere with both RNAi and interferon as these two systems are dependent on the availability of viral RNA substrates (Li et al., 2004).

Given the data presented, the emergence of negative-sense RNA viruses would seem counterintuitive. Based on recent evolutionary predictions, the current hypothesis concerning the origins of this phylum derive from the rapid expansion of positive-sense RNA viruses which branched off into double stranded RNA viruses thereby providing an opportunity to utilize the negative-sense strand in isolation (Wolf et al., 2018). However, utilizing an RNA genome with negative polarity poses a number of inherent constraints. The most prominent of these is the inherent immunogenicity of the viral uncapped RNA that is reliant on the interaction of RdRp or extensive secondary structure to protect its 5’ end, together with the maintenance of high levels of NP to shield its genome (Gebhardt et al., 2017). As shown herein, small perturbations to the balance of NP availability showcases the fragility of this system and its general lack of robustness. Perhaps it is for this reason that negative-sense RNA viruses show the lowest diversity within the known global virosphere (Walker et al., 2019; Wolf et al., 2018).

The dynamics of negative-sense RNA viruses and their inherent biological conflict between replication and host recognition demand the production of potent antagonists in order to enhance their adaptability despite their constrained fitness. This is exemplified in our transcriptomic data comparing wild type IAV to viruses in which NS1 is lacking or limited. While NS1, like many viral antagonists regardless of virus group, inhibits the direct cellular antiviral response, our results show that NS1 interferes with an expansive range of cellular processes including: immune cell recruitment, cell death, development, and metabolism. Similar results have been accredited to VP35 – the RNA binding viral antagonist of Ebola virus (Albarino et al., 2018; Cardenas et al., 2006).

Regardless of the potency of a given viral antagonist, our data show that PAMP production can rapidly overwhelm the capacity of the virus to mask it and lead to induction of the antiviral defenses. To minimize this occurrence, NP provides a scaffold for the genome to enable full-length replication and transcription by the viral polymerase upon entry. This biology generally allows for initial virus transcription to occur in the absence of interferon. As replication begins, nascent genomes demand increasing amounts of NP, which, if unmet, result in the production of DVGs. It is during this inevitable phase where antagonists such as NS1 of IAV, P frameshifts of paramyxoviruses, or the M protein of rhabdoviruses that are required to further buffer the host response, allowing the virus to complete its life cycle (Garcia-Sastre, 2017).

Because of its critical nature, NP is also under significant structural constraints to bind RNA and support RdRp function (Reguera et al., 2014). These constraints, together with the inherent propensity for DVG formation in conditions of limited NP, make this protein an ideal drug target. What is particular attractive about this concept is the potential for bystander effect as NP targeting, even in a subset of cells, could trigger the amplification of both DVGs and the host interferon response leading to replication catastrophe and thereby serving as an effective and universal antiviral therapeutic.

## MATERIALS AND METHODS

### Cell Culture

A549, A549-STAT1^-/-^, HEK293T, HEK293T-NoDice, HeLa, Vero, Vero-E6, BHK-J, MDCK, and MDCK-NS1-GFP cells were maintained at 37°C and 5% CO_2_ in Dulbecco’s Modified Eagle Medium (DMEM, Gibco®) supplemented with 10% Fetal Bovine Serum (FBS, Corning®) as described elsewhere (Bogerd et al., 2014; Chua et al., 2013). Normal human bronchial epithelial (NHBE) cells (Lonza, CC-2540 Lot# 580580) were isolated from a 79-year-old Caucasian female and were maintained in bronchial epithelial growth media (Lonza, CC-3171) supplemented with BEGM SingleQuots as per the manufacturer’s instructions (Lonza, CC-4175) at 37°C and 5% CO_2_.

### Viruses

Influenza A/Puerto Rico/8/34 (H1N1) viruses (NCBI:txid183764) with low (LD) and high (HD) DVG content were grown for 48 h in MDCK cells (for wild type viruses) or MDCK-NS1 cells (for ΔNS1 viruses (Garcia-Sastre et al., 1998)) at an MOI of 0.001 (LD) or 0.05 (HD) in DMEM supplemented with 0.3% BSA (MP Biomedicals®) and 1 μg/ml TPCK-trypsin (Sigma-Aldrich®). The IAV-NS1-T virus was derived from influenza A/Puerto Rico/8/34 (H1N1) virus and was constructed following a previously described strategy where tandem target sites for miRNA-30 were inserted into the 3′ UTR of the NS1 gene (Chua et al., 2013). mNeon-expressing influenza A/Puerto Rico/8/34 (H1N1) virus (IAV-mNeon) was a kind gift from Peter Palese. Infectious titers of influenza A viruses were determined by plaque assays in MDCK or MDCK-NS1 cells. Influenza A/Puerto Rico/8/34 (H1N1) virus and Sendai virus containing a miRNA-silencing cassette targeting viral NP transcripts (IAV-NP-T and SeV-NP-T, respectively) as well as their respective control viruses (IAV-NP-C and SeV-NP-C) were rescued as previously described (Aguado et al., 2018). IAV-NP-T, IAV-NP-C, and SeV-NP-C were grown in 10-day old specific pathogen-free (SPF) chicken eggs (Charles River Laboratories). SeV-NP-T virus was grown on HEK293T cells lacking hDicer functionality (HEK293T-NoDice) for 3-5 days. Viral titers of IAV-NP-T were determined by egg infectious dose (Aguado et al., 2018). SeV-NP-T virus was titered on NoDice 293T cells by TCID50.Infections with wild-type or mutant IAV were performed at the indicated MOIs for 1 h at room temperature in DMEM supplemented with 0.3 % BSA and 1 μg/ml TPCK-trypsin before incubation for the indicated amount of time at 37°C. For infection of NHBE cells with IAV, cells were washed with PBS after initial adsorption of the virus and supplemented with fresh infection media. Infections with SeV-NP-T/C were performed at the indicated MOIs in DMEM supplemented with 10% FBS and incubated for the indicated amount of time at 37°C. Recombinant trisegmented Lassa virus expressing tdTomato and *Renilla* luciferase (rLASV-tdTom) was generated using a previously successful strategy to create a trisegmented recombinant lymphocytic choriomeningitis virus expressing two additional genes of interest (Emonet et al., 2009) and was based on LASV strain Bantou 366 (Ba366) which was obtained from the Institute of Virology at the University of Marburg (Lecompte et al., 2006). The coding capacity of this recombinant virus was extended by incorporation of two S segments instead of one, one of these carrying the NP and *Renilla* luciferase genes and the other carrying the GPC and tdTomato genes. The plasmid pHH21-LASV-Sag (GenBank #GU830839.1) was used to generate the pHH21-LASV-Sag-GPC/tdTomato and pHH21-LASV-Sag-*Renilla*/NP constructs. For confirmatory purposes of generated viruses later on, silent mutational markers were introduced into the NP (A1252T) and GPC (G377A) genes at the nucleotide level. The tdTomato gene (EU855182.1) carries the marker mutation A1244G and was cloned from the tdTomato-pBAD plasmid (a kind gift from Drs. M. Davidon, N. Shaner, and R. Tsien, Addgene plasmid #54856). The humanised reporter gene *Renilla* luciferase (GenBank AF362549) was cloned from the previously described LASV Ba366 minigenome reporter plasmid (Oestereich et al., 2016). Approximately 4 × 10^5^ BHK-J cells were co-transfected with 0.75 µg of pCAGGS-LASV-NP (GenBank # GU830839.1; ADI39452.1), 1.5 µg of pCAGGS-LASV-L (GenBank #GU979513.1; ADU56645.1), 1.5 µg of pHH21-LASV-Lag (GenBank #GU979513.1), 0.75 µg of pHH21-LASV-Sag-GPC/tdTomato and 0.75 µg of pHH21-LASV-Sag-Renilla/NP using Lipofectamine 2000 according to the manufacturer’s instructions. At 4 h post-transfection, media was replaced with DMEM supplemented with 5% FBS. One day post-transfection, cells and supernatants were transferred to a T75 flask. At 5 days post-transfection, cells and supernatants were transferred to a T175 flask. Every 3 to 4 days, supernatants were harvested, infected cells were passaged and 8 ml of supernatant with 15 ml fresh medium was added to the cells. Cells were tested for viral infection at each passage by immunofluorescence staining and viral titers in supernatants were determined by immunofocus assays as described elsewhere (Rieger et al., 2013). The correct sequence of the recombinant virus was confirmed by sequencing. Recombinant GFP-expressing Ebola virus/H.sapiens-tc/COD/1976/Yambuku-Mayinga (rgEBOV-GFP, GenBank KF990213.1, henceforth called EBOV-GFP) was previously described(Hoenen et al., 2013). EBOV-eGFP was grown in Vero-E6 cells and viral titers were determined by immunofocus assays as described previously (Escudero-Perez et al., 2019). Rescue of eGFP- and GLuc-expressing Human parainfluenza virus 3, strain JS (rHPIV3^JS^-GlucP2AeGFP, henceforth called HPIV3-GFP) was carried out at 32°C as previously described (Beaty et al., 2017). For virus amplification, rescue supernatant was transferred to HeLa cells at 32°C in DMEM supplemented with 10% FBS. Once >90% of cells were eGFP-positive, culture medium was replaced with DMEM supplemented with 1ug/mL TPCK-trypsin (Millipore Sigma, Burlington MA, USA) for 24h at 32°C. Thereafter, supernatant was collected and clarified of cell debris by centrifugation. HPIV3-eGFP was titered on Vero cells by serial dilution, and infectious units were defined by GFP-positive events at 24h post-inoculation, reported as iu/mL. Rescue of recombinant eGFP-expressing measles virus, strain Edmonston B (rMeV^EdmonstonB^-eGFP, henceforth called MeV-GFP) was carried out as described previously^23^. MeV-eGFP was amplified in Vero cells from an MOI of 0.01. 12 h after cells were 100% infected (as defined by visual determination of GFP positive cells), cells were collected into the supernatant and pipetted vigorously to liberate cell-associated virus. Cellular debris was then cleared by centrifugation, and virus stocks were titered as described above for HPIV3-GFP. Recombinant GFP-expressing human respiratory syncytial virus, strain A2 (rgRSV[224], henceforth called HRSV-GFP) was generously provided by Dr. M. Peeples (OSU) and was described previously (Hallak et al., 2000). Recombinant Indiana Vesicular Stomatitis Virus expressing eGFP (VSV-GFP) was generated by inserting an eGFP open reading frame in the intergenic region between the M and G open reading frames using the reverse genetics system provided by Dr. G. Wertz (UVA) and described elsewhere (Whelan et al., 1995).

### Western Blot

Cells were lysed in NP40 lysis buffer containing 1× cOmplete Protease Inhibitor Cocktail (Roche®) and 1× Phenylmethylsulfonyl fluoride (Sigma Aldrich®) and cleared from the insoluble fraction by centrifugation at 17,000 × *g* for 5 min at 4°C. For cells infected with EBOV-GFP and LASV-tdTom, cells were lysed in SDS sample buffer and samples were inactivated for 10 min at 95°C prior to transfer of samples out of the BSL-4 laboratory. Samples were separated by SDS-PAGE and transferred onto nitrocellulose membranes. Proteins were detected using mouse monoclonal anti-Actin (Thermo Scientific, MS-1295), rabbit monoclonal anti-IFIT1 (Cell Signalling, D2X9Z), rabbit polyclonal anti-GFP (Abcam, ab290), mouse monoclonal anti-IAV NP (Center for Therapeutic Antibody Discovery at the Icahn School of Medicine at Mount Sinai, clone HT103), mouse monoclonal anti-IAV NS1 (Center for Therapeutic Antibody Discovery at the Icahn School of Medicine at Mount Sinai, clone 1A7), mouse monoclonal anti-IAV M1/M2 (Center for Therapeutic Antibody Discovery at the Icahn School of Medicine at Mount Sinai, clone E10), mouse monoclonal anti-SeV NP (Center for Therapeutic Antibody Discovery at the Icahn School of Medicine at Mount Sinai, clone 6H4), rabbit polyclonal anti-HPIV3 NP (GenScript, custom made against peptide: CNINSSETSFHKPTG),), mouse monoclonal anti-EBOV NP (Invitrogen, MA5-29991) and previously described (Amanat et al., 2018) mouse monoclonal anti-LASV GP (generously provided by Dr. F. Krammer, KL-AV-1B3) primary antibodies. Primary antibodies were detected using HRP-conjugated secondary anti-mouse (GE Healthcare®, NA931V) and anti-rabbit (GE Healthcare®, NA934V) antibodies and visualized using Immobilon Western Chemiluminescent HRP substrate kit (Millipore®) according to the manufacturer’s instructions.

### RNA sequencing

**T**otal RNA from infected and mock treated cells was extracted using TRIzol (Thermo Fisher®) or the RNeasy Mini kit (QIAGEN®) according to the manufacturer’s instructions and treated with DNase I. RNA-seq libraries were prepared using the TruSeq RNA Library Prep Kit v2 (Illumina®) for mRNA and TruSeq Stranded Total RNA Library Prep Gold (illumina) for total RNA following manufacturer’s instructions. All sequencing runs were performed using an Illumina NextSeq 500 platform.

### Analysis of sequencing data

Sequencing reads were aligned to the human genome (hg19) using the STAR aligner (Dobin et al., 2013) followed by differential gene expression analysis by DESeq2 (Love et al., 2014). Heatmap plots of statistically significant DEGs (L2FC ≥ 2, Adjusted p-value < 0.05) were constructed using the heatmap.2 function form the gplots package (https://CRAN.R-project.org/package=gplots) in R (http://www.R-project.org/). Diffusion maps were constructed with the destiny package (Angerer et al., 2016) in R, using the logarithm base 2 of the transcript per million (TPM) values for statistically significant DEGs in at least one experimental condition. Sequencing reads were also aligned to respective viral genomes using Bowtie2 (Langmead and Salzberg, 2012) and visualized using IGV software (Thorvaldsdottir et al., 2013). The viral genome references used were SeV: KY295909.1 and IAV (A/Puerto Rico/8/34/Mount Sinai): AF389122.1, AF389121.1, AF389120.1, AF389119.1, AF389118.1, AF389117.1, AF389116.1, AF389115.1. Defective viral genomes (DVGs) identification and quantification was performed using ViReMa (Routh and Johnson, 2014) following a previously described pipeline (Alnaji et al., 2019). Only strand congruent recombination events were quantified for IAV (to account for deletion DVGs), whereas all viral recombination events were quantified for SeV (to account also for copy back DVGs). DVG junction coordinates were visualized by sushi plots using the sushi package (Phanstiel et al., 2014) in R. miRNA expression in NHBE cells was identified using the Small RNA application (Illumina®, BaseSpace) on previously published NHBE small RNA sequencing data (SRA: SRR5127216) (McCall et al., 2017). Sequencing data generated in this manuscript is available on NCBI GEO under accession number GSE122794.

### Gene set enrichment analysis

In order to select genes whose expression was differentially affected by IAV NS1 levels, we first ranked each gene based on their expression (TPM; transcripts per million) values i) in cells infected with IAV-ΔNS1 versus cells infected with IAV-WT or IAV-30T (genes responsive to the presence of any NS1), or ii) in cells infected with IAV-WT versus cells infected with IAV-ΔNS1 or IAV-NS1-T (genes responsive to WT NS1), using Gene Set Enrichment Analysis (GSEA) (Subramanian et al., 2005). The sign of the difference between the rank metric scores (Signal to Noise ratio) of both comparisons (any NS1 - WT NS1) defined whether the gene was responsive to any NS1 (negative) or only to WT NS1 levels (positive). Only genes that had a rank metric score < -1 and were statistically significantly differentially expressed (in any comparison or condition) were taken into account. Finally, the membership to a specific pattern (genes responsive to any NS1 or WT NS1) was weighted taking into account the fold change in expression (L2FC; log2 fold-change) of cells infected with IAV-WT (WT NS1) or IAV-NS1-T compared to IAV- ΔNS1 infections. The weighted list of selected genes for each pattern was used to identify enriched gene ontology (GO) annotations (biological process) using Enrichr (Kuleshov et al., 2016). Redundant GO annotations were reduced by eliminating annotations that had > 75% overlap to another GO annotation using the reduce_overlap function from the GOplot package in R (Walker et al., 2019). Final annotations were visualized in R using custom scripts.

### RNP reconstitutions

Approximately, 1 × 10^6^ HEK-293T cells were transiently transfected with 1 μg each of pcDNA-PB2 (Fodor et al., 2002), pcDNA-PB1 (Fodor et al., 2002) or pcDNA-PB1a (Vreede et al., 2004) and pcDNA-PA (Fodor et al., 2002), 4 μg of pPOLI-vNA200 and the indicated amounts of pcDNA-NP (Fodor et al., 2002) using Lipofectamine 2000 and Opti-MEM according to the manufacturer’s instructions. The empty vector pcDNA-3a was used to balance total amounts of transfected DNA. The plasmid pPOLI-vNA200 was derived from the previously described plasmid pPOLI-NA (Fodor et al., 1999) and constructed using PCR-mediated deletion mutagenesis. Cells were harvested 48 h post-transfection and total RNA was extracted using TRIzol (Thermo Fisher®) according to the manufacturer’s instructions.

### Northern Blot

Detection of viral RNA by Northern blot was performed in a similar manner as previously described with the following modifications (Perez et al., 2010). Briefly, 30 μg of total RNA was resolved by 12% 7M urea PAGE in TBE buffer and transferred onto Hybond NX membranes (Amersham®) using an Owl HEP-1 Semi Dry Electroblotting System (Thermo Scientific®). RNA was chemically crosslinked with EDC cross-linking solution (0.16 M 1-ethyl-3-(3-dimethylaminopropyl) carbodiimide, 0.13 M 1-methylimidazole, pH = 8.0) for 1 h at 65°C, prior to blocking in 6× SSC and 7% SDS for 1 h at 65°C. Membranes were hybridized with radiolabeled probes against 5′ vRNA termini (5′-AAAAANNNCCTTGTTTCTACT-3′) and U6 snRNA (5′-GCCATGCTAATCTTCTCTGTATC-3′) at 30°C for 12 h. Membranes were washed three times with 3× SSC and 0.1 % SDS) at 30°C for 15 min. Radiolabeled probes were detected by autoradiography using a Typhoon Trio Variable Mode Imager (GE Healthcare®).

### RT-PCR analysis

For qualitative analysis of viral and cellular RNA, total RNA was extracted using TRIzol (Thermo Fisher®) or the RNeasy Mini kit (QIAGEN®) and treated with DNase I according to the manufacturer’s instructions. Extracted RNA was reverse transcribed using SuperScript II and oligo (dT) primers or universal IAV primers (Table S10). cDNA from viral RNA or cellular transcripts were amplified using primers listed in Table S10 and GoTaq Green MasterMix (Promega®). PCR products were analysed by 1.5% agarose gel electrophoresis in TAE buffer.

### Chemical inhibition of IAV

To determine the inhibitory role of different compounds on viral replication, A549 cells were infected for 1 h at room temperature with influenza A/Puerto Rico/8/34 (H1N1) virus at an MOI of 5 in DMEM supplemented with 0.3% BSA. 1 h post-infection the indicated amounts of thapsigargin (Sigma-Aldrich®, 586006), nucleozin (Calbiochem®, 492905) or Baloxavir marboxil (eNovation Chemicals®, D621084), reconstituted in DMSO, were. added using DMSO as a control Cells were incubated at 37°C for 8 h before being harvested as described before and analyzed by Western blotting.

### RNAi-mediated silencing of viral nucleoprotein expression

Custom siRNA pools (Custom siGENOME SMARTpool, Dharmacon®) were designed against the nucleoprotein of influenza A/Puerto Rico/8/34 (PR8) virus (GenBank: AF389119.1), human respiratory syncytial virus strain A2 (GenBank: KT992094.1), human parainfluenza virus 3 strain JS (GenBank: Z11575.1), measles virus strain Edmonston (GenBank: DQ839356.1), Ebola virus/H.sapiens-tc/COD/1976/Yambuku-Mayinga (NCBI Reference Sequence NC_002549.1) and Lassa mammarena virus strain BA366 (GenBank: GU830839.1). siRNA sequences are detailed in Table S10. Approximately 1 × 10^5^ A549 cells were transfected with 20 pmoles Control siRNA-A (Santa Cruz Biotechnology®, sc-37007) or NP specific siRNA using Lipofectamine RNAiMax and Opti-MEM according to the manufacturer’s instructions and incubated in DMEM supplemented with 10% FBS for 24 h at 37°C. Prior to infection, cell monolayers were washed with PBS. Infections with IAV-mNeon were performed at an MOI of 1.5 for 1h at room temperature in DMEM supplemented with 0.3% BSA and 1 μg/ml TPCK-trypsin prior to incubation for 12 h at 37°C. Infections with HRSV-GFP were performed at an MOI of 5 for 1 h at room temperature in DMEM supplemented with 0.3% BSA prior to incubation for 12 h at 37°C. Infections with HPIV3-GFP were performed at an MOI of 5 for 1 h at room temperature in DMEM supplemented with 0.3% BSA prior to incubation for 12 h at 37°C. Infections with VSV-GFP were performed at an MOI of 5 for 6 h at 37°C in DMEM supplemented with 10% FBS. Infections with MeV-GFP were performed at an MOI of 0.1 for 48 h at 37°C in DMEM supplemented with 10% FBS. Infections with EBOV-GFP were performed at an MOI of 1 in DMEM for 1 h at 37°C, before removal of the viral inoculum and incubation of infected cells in DMEM supplemented with 3% FBS for 16 h at 37°C. Infections with rLASV-tdTom were performed at an MOI of 1 in DMEM for 4 h at 37°C, before removal of the viral inoculum and incubation of infected cells in DMEM supplemented with 10% FBS for 20 h at 37°C.At the indicated time points, cells were harvested using trypsin and analysed by flow cytometry and Western blot as described above.

### Flow cytometry

Cells infected with IAV-mNeon, HRSV-GFP, HPIV3-GFP and VSV-GFP were fixed and inactivated in 4% formaldehyde for 30 min at room temperature. Fixed cells were diluted in PBS to 500-1000 cells/μl followed by analysis by FACS on a Guava EasyCyte flow cytometer (Millipore®). EBOV-GFP infected cells were fixed and inactivated in 4% formaldehyde for 30 min at room temperature prior to four washed in PBS. FACS data was acquired on a LSRFortessa (BD Biosciences®). rLASV-tdTom infected cells were inactivated and fixed in 4% formaldehyde for 30 min at room temperature prior to four washes in PBS. Fixed cells were resuspended in PBS supplemented with 1% FBS and 1 mM EDTA and filtered through a cell strainer. 10,000 cells per sample were analyzed by FACS on a FACSAria III cell sorter (BD Biosciences, 531 nm laser, BP 585/15). Acquired FACS data was analyzed using FlowJo to quantify fluorescent reporter gene expressing cells in each sample set.

### Biosafety

All viral experiments were performed under proper biocontainment. IAV, MeV, SeV, HPIV3, VSV, and HRSV were performed under BSL2+ conditions at the Icahn School of Medicine at Mount Sinai. Rescue and evaluation of recombinant EBOV or LASV were performed under BSL-4 conditions in the BSL-4 laboratory at the Bernhard Nocht Institute for Tropical Medicine (Hamburg, Germany).

## ACKNOWLEDGEMENTS

We wish to thank Dr. Lisa Oestereich and Nadja Höfs for providing the plasmids used for cloning rLASV-tdTom; Dr. Florian Krammer for the LASV-GP antibody. This work was supported by the INTERfering and CO-Evolving Prevention and Therapy (INTERCEPT) program sponsored by DARPA (DARPA-16-35-INTERCEPT-FP-006). D.B-M is an Open Philanthropy Fellow of the Life Sciences Research Foundation (LSRF). P.A.T. was supported by a Canadian Institutes of Health Research (CIHR) Postdoctoral Fellowship. B.L was supported by NIH grant AI115226. S.O. was supported by the Jürgen Manchot foundation Ph.D. fellowship.

## FIGURES AND FIGURE LEGENDS

**Figure S1:**
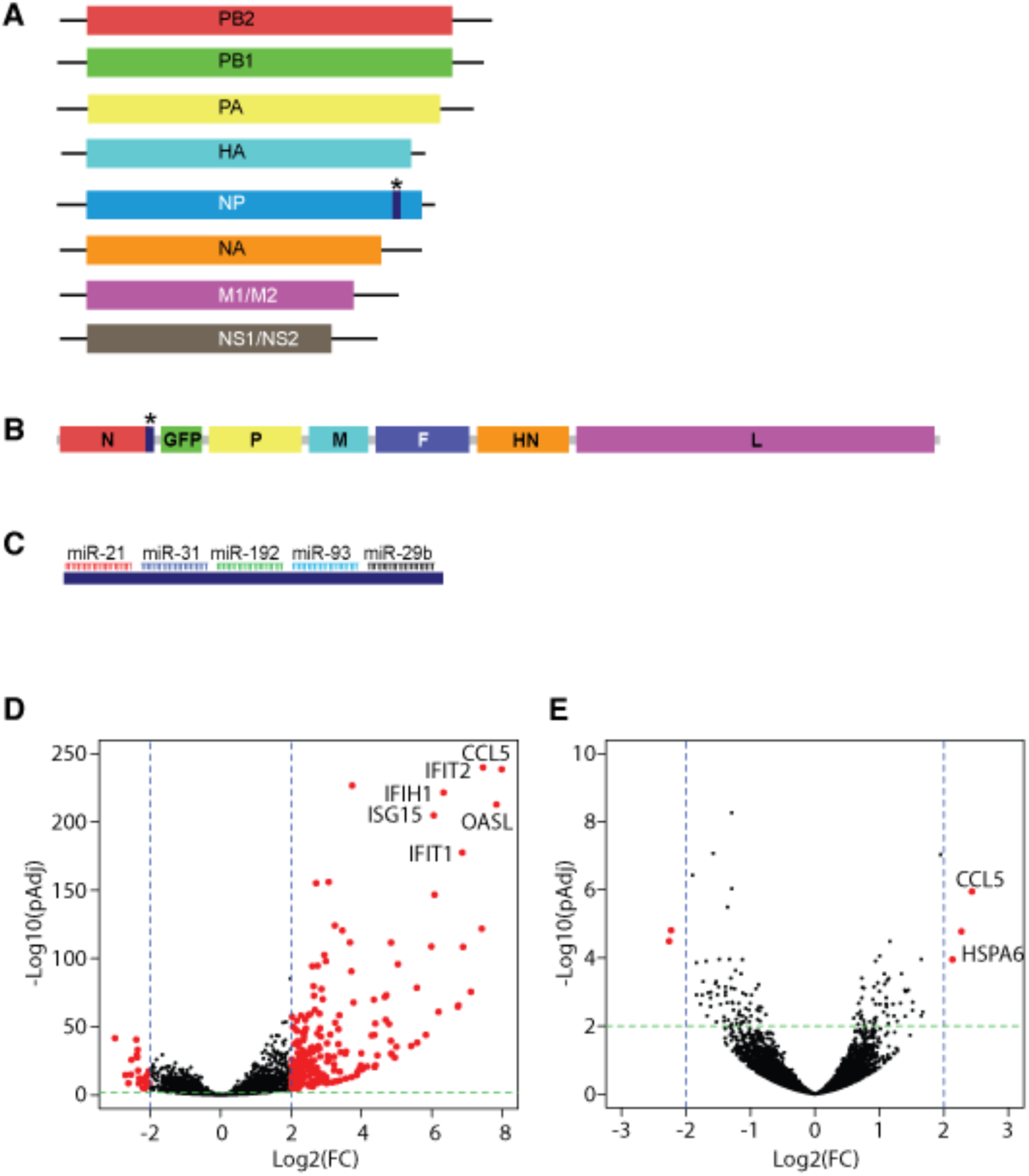
Strategy for NP targeting and transcriptional response of non-targeted viruses. (**A**-**C**), Diagram of recombinant (**A**) IAV-NP-T and (**B**) SeV-NP-T where the (**C**) miRNA targeting casette is downstream of the nucleoprotein (NP/N) ORF. SeV-NP-T also has a GFP ORF inserted between N and the phosphoprotein. **D**-**E**, Volcano plots depicting the host transcriptional response to (**D**) SeV-NP-C and (**E**) IAV-NP-C. Red points represent statistically significant differentially expressed genes (DEGs) in A549 cells infected with the corresponding viruses compared to mock infected cells (Log2(FC) > 2, pAdj < 0.01, n=2).

**Figure S2:**
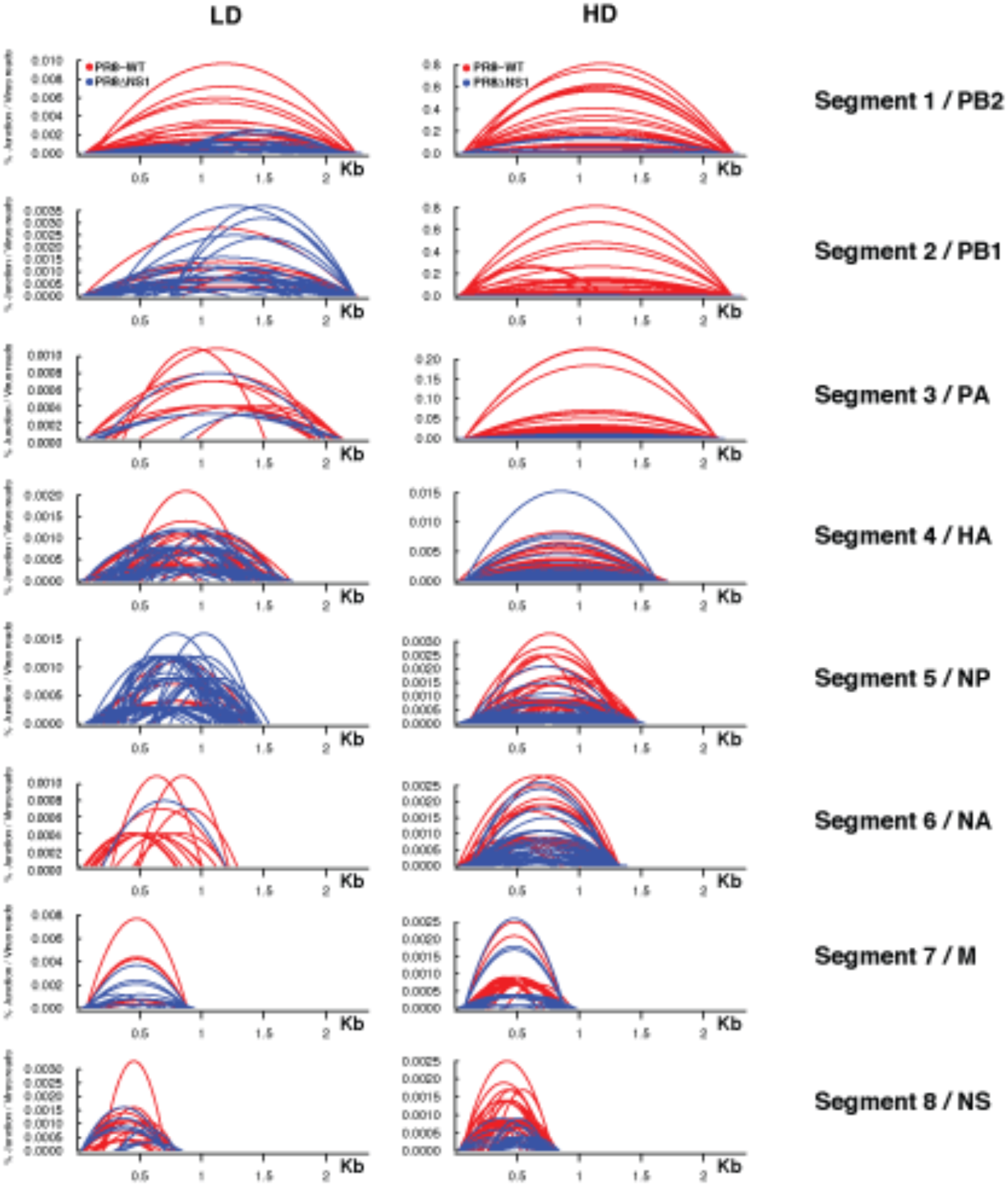
Segment-specific defective viral genome species. Analysis of mRNA-seq reads aligning to the viral genome for DVGs present in NHBE cells infected with LD or HD versions of wild-type IAV or IAV-ΔNS1 virus (MOI = 3, 12 hpi). Junctions sites of DVGs were visualised across each viral segment. x-axis denotes nucleotide position and y-axis corresponds to the percentage of junction reads over total number of viral reads (n = 3).

**Figure S3:**
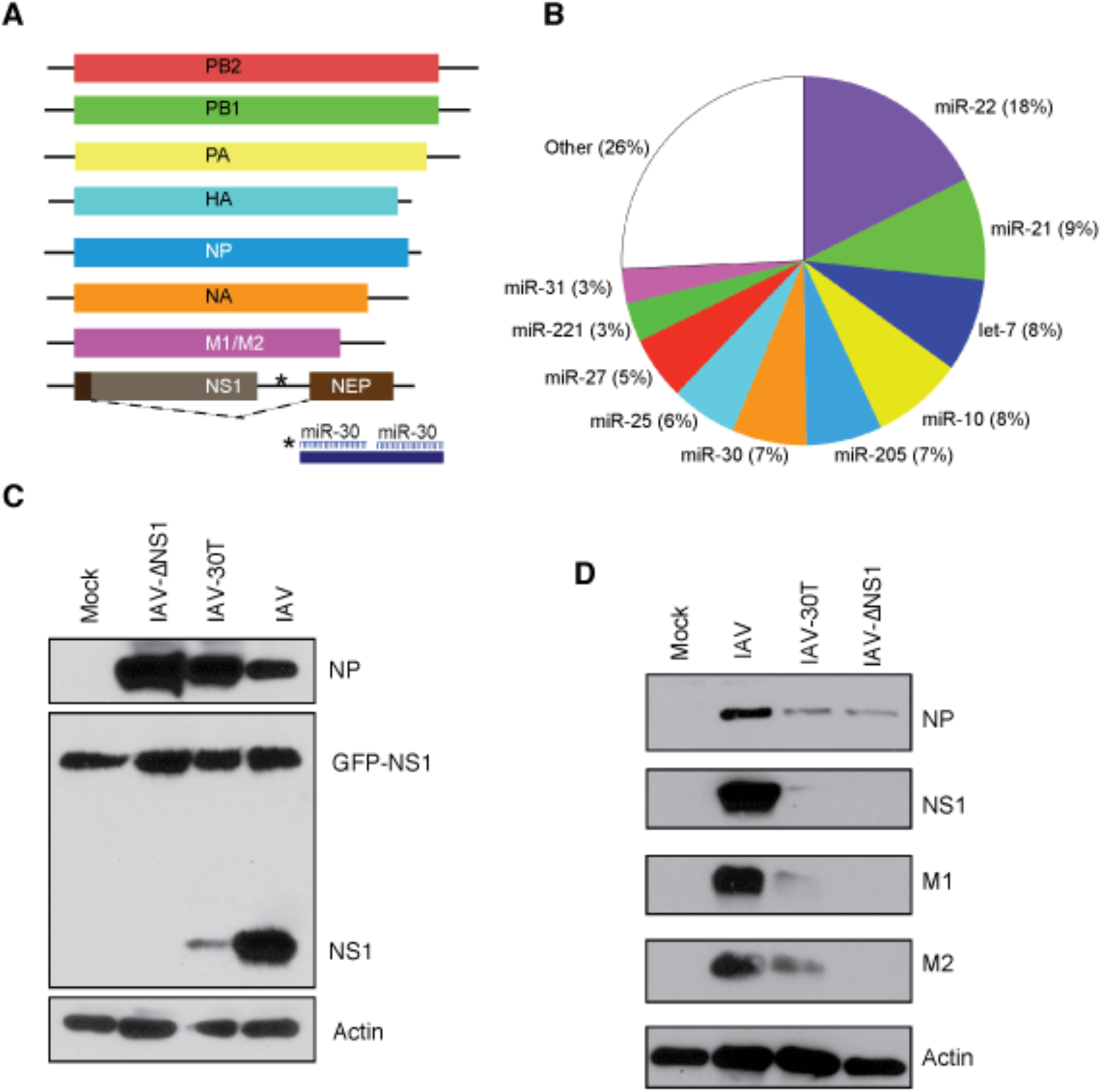
Modulation of IAV NS1 protein expression by miRNA targeting. **(A**) Diagram of recombinant IAV-NS1-T where two miR-30 target sites were introduced into the 3’ UTR of the NS1 ORF. **(B)** miRNA expression profile of NHBE cells at baseline. Values represent the percentage of total miRNA reads for each miRNA. Data from (McCall et al., 2017). (**C**) MDCK-NS1 cells, constitutively expressing a NS1-GFP fusion protein were infected with wild-type IAV, IAV-ΔNS1 or IAV-NS1-T for 12 hrs at an MOI of 1. Whole cells extracts were analysed by Western blot for viral protein expression. (**D**) NHBE cells were infected with IAV-WT, IAV-ΔNS1 or IAV-NS1-T for 12 h at an MOI of 3. Whole cells extracts were analysed by Western blot for viral protein expression.

**Figure S4:**
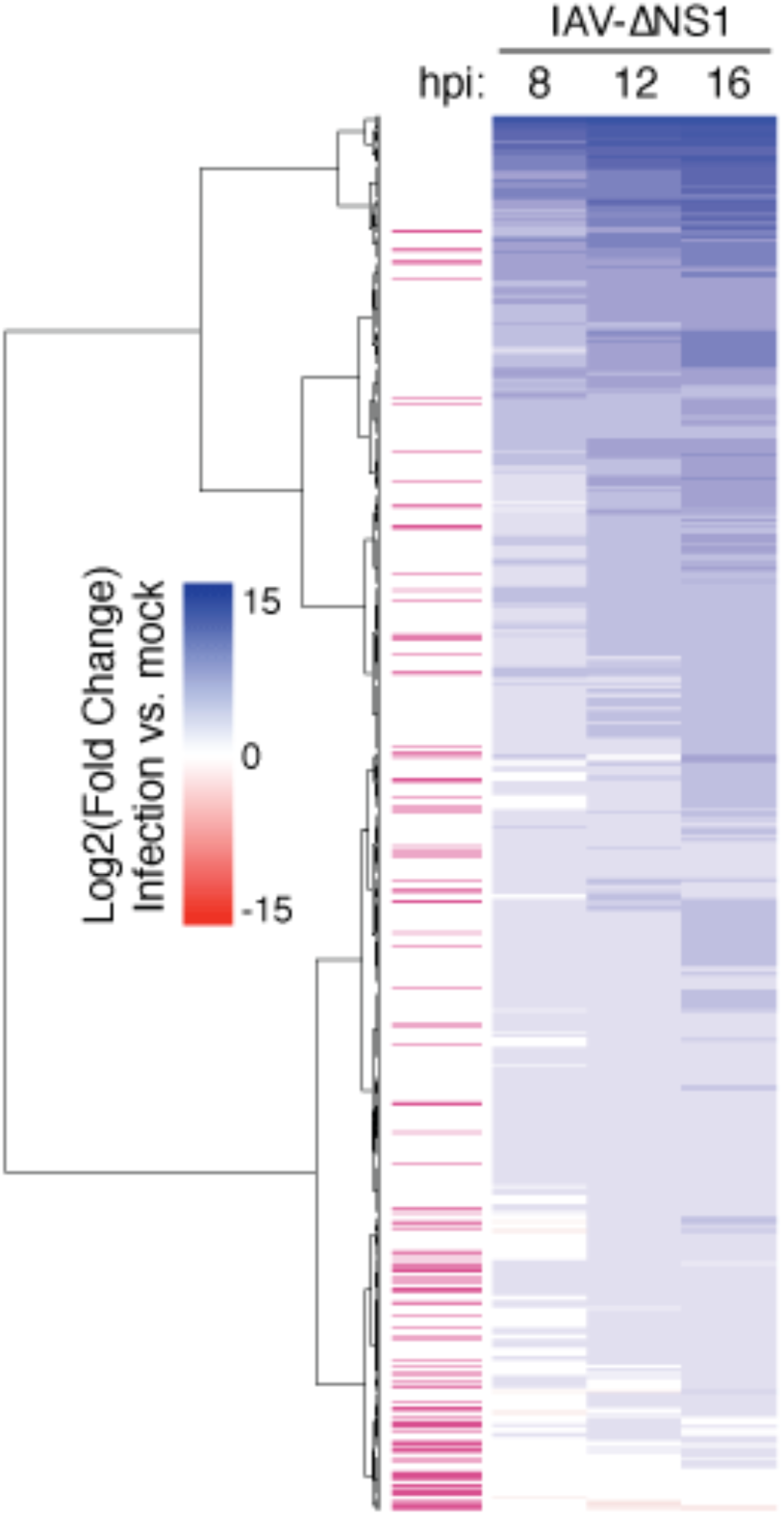
Modulating the expression of NS1 reveal dynamics of transcriptional inhibition. Heatmap depicting the transcriptional response of NHBE cells infected with IAV-ΔNS1 for 8, 12 and 16hrs (n = 2, MOI = 3). Values represent the Log_2_(fold change) expression (compared to mock-treated cells) of differentially expressed gene markers for the NS1 negative vs. NS1 positive IAVs comparison (colored in white), or the WT vs. mutant NS1 IAVs comparison (colored in magenta).

**Figure S5:**
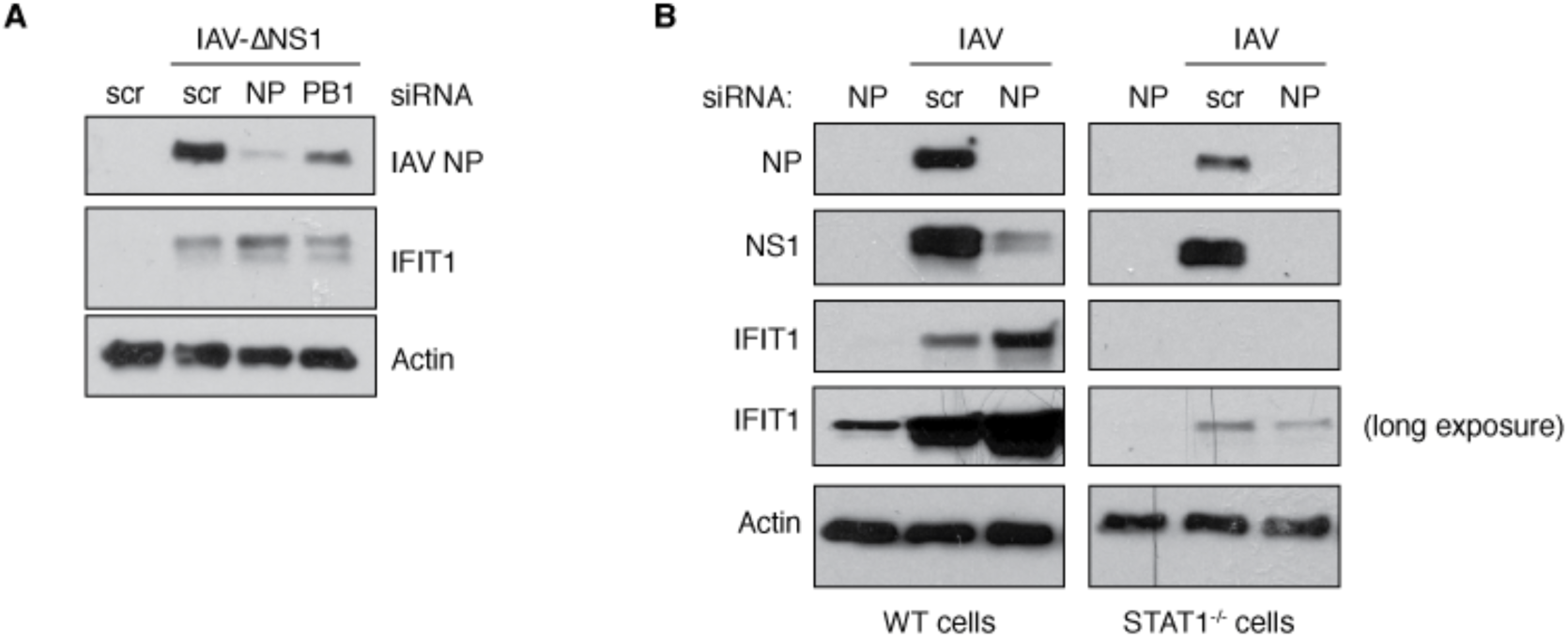
Determining how loss of NP vs. PB1 impacts IFN-mediated IFIT1 levels. **A**, A549 cells were transfected with siRNA targeting IAV NP or PB1 transcripts or a scrambled control siRNA prior to infection with IAV-ΔNS1 at an MOI of 2 for 8 h. Whole cells extracts were analysed by Western blot for viral protein expression. **B**, Wild type or STAT^-/-^ A549 cells were transfected with siRNA targeting IAV NP or a scrambled control siRNA prior to infection with wild type IAV at an MOI of 5 for 8 h. Whole cells extracts were analysed by Western blot for viral protein expression.

**Figure S6:**
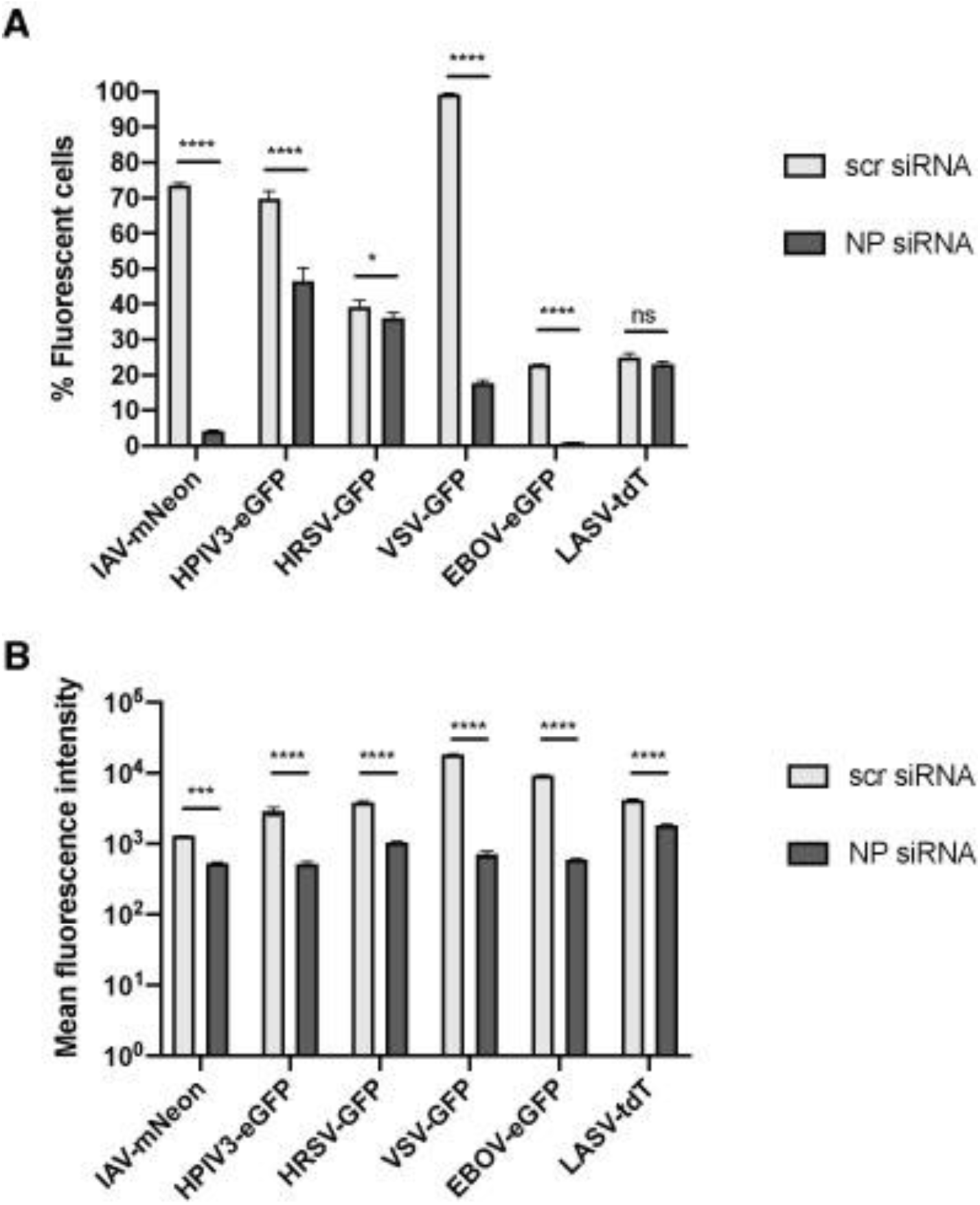
siRNA-mediated inhibition of negative-sense RNA virus infections. **(A)** A549 cells were transfected with siRNA targeting virus-specific NP transcripts prior to infection with a wide range of negative-sense RNA viruses as described in Figure 6. Fixed cells were analysed by flow cytometry for virus-induced fluorescent reporter gene expression. The graph shows the mean percentage of fluorescent positive cells from three biological replicates. Error bars represent standard deviations. (**b**) Mean fluorescence intensity of each cell analysed in (**A**). The graph shows the mean fluorescence intensity from three biological replicates. Significance was determined by two-tailed two sample *t* tests: ns, * p < 0.05, *** p < 0.001, **** p < 0.0001.

## Supplemental Tables 1-10

All supplemental tables and legends can be downloaded using the following link: https://bit.ly/383k2BV

